# PhenoVision: A framework for automating and delivering research-ready plant phenology data from field images

**DOI:** 10.1101/2024.10.10.617505

**Authors:** Russell Dinnage, Erin Grady, Nevyn Neal, Jonn Deck, Ellen Denny, Ramona Walls, Carrie Seltzer, Robert Guralnick, Daijiang Li

## Abstract

Plant phenology plays a fundamental role in shaping ecosystems, and global change-induced shifts in phenology have cascading impacts on species interactions and ecosystem structure and function. Detailed, high-quality observations of when plants undergo seasonal transitions such as leaf-out, flowering, and fruiting are critical for tracking causes and consequences of phenology shifts, but these data are often sparse and biased globally. These data gaps limit broader generalizations and forecasting improvements in the face of continuing disturbance. One solution to closing such gaps is to document phenology on field images taken by public participants. iNaturalist, in particular, provides global scale research-grade data and is expanding rapidly. Here we utilize over 53 million field images of plants and millions of human annotations from iNaturalist – data spanning all angiosperms and drawn from across the globe – to train a computer vision model (PhenoVision) to detect the presence of fruits and flowers. PhenoVision utilizes a vision transformer architecture pretrained with a masked autoencoder to improve classification success, and it achieves high accuracy for flower (98.5%) and fruit presence (95%). Key to producing research-ready phenology data is post-calibration tuning and validation focused on reducing noise inherent in field photographs, and maximizing the true positive rate. We also develop a standardized set of quality metrics and metadata so that results can be used effectively by the community. Finally, we showcase how this effort vastly increases phenology data coverage, including regions of the globe where data have been limited before. Our end products are tuned models, new data resources, and an application streamlining discovery and use of those data for the broader research and management community. We close by discussing next steps, including automating phenology annotations, adding new phenology targets, e.g., leaf phenology, and further integration with other resources to form a global central database integrating all in-situ plant phenology resources.

## Introduction

Phenology – the timing of life-cycle events such as leaf budburst, flowering, and fruiting – is highly responsive to environmental conditions and plays a fundamental role in shaping ecosystems. For millennia, phenology has shaped human survival, and formalized efforts to document the timing of seasonal events have been in place worldwide for centuries, mainly to support agricultural practices [1,2]. These long-term records have been crucial for understanding patterns and drivers of phenological transitions [3–6]. Changes to phenology, triggered by shifting environmental conditions, can have cascading impacts on multi-species population dynamics, species distributions, trophic interactions,and ecosystem services such as carbon dynamics, water and nutrient cycling, and albedo [7–9]. These changes can trigger dramatic, and sometimes devastating, consequences for ecosystems, with negative implications for human health and economic interests [3,6].

Despite the importance of phenology for global ecosystem structure and function, large phenology data gaps exist [10]. As a result, phenology studies are often narrow in their spatial, temporal, and taxonomic breadth, limiting broader, macrosystems-level insights into patterns and drivers of phenological change [8,10–12]. Researchers increasingly recognize the importance of large-scale phenological analysis covering broad geographic areas and climatic gradients, various habitat types and land use categories, and many taxa. Such approaches, leveraging growing data resources and flexible modeling frameworks, can identify emergent signals of change and how abiotic and biotic drivers interact with plant physiological mechanisms to shape species and community phenology responses [13,14]. While recent studies have showcased the power of mature modeling frameworks and growing data sources, two key challenges are closing global phenology data gaps and developing better integration of disparate resources. Developing a consistent, global view of plant phenology from canopy to understory and across key phenological phases for leaves, flowers, and fruits requires new approaches for assembling and serving high quality, research-ready and interoperable data.

One area of potential transformative value for phenology is using community-collected field images that have been vetted in terms of their identification. In particular, the iNaturalist platform, which is producing millions of plant photographs per month as of 2024 and is still accelerating in terms of user involvement, may be uniquely poised to fill critical gaps in global phenology data. The iNaturalist platform allows users to post an image of any organism found in its environment, along with occurrence metadata and a preliminary identification [15]. A majority-rule based community identification process provides a validated taxonomic identification to the lowest possible taxonomic unit, most often species [16]. While such data resources are biased in terms of sampling [17], they capture phenological phases across diverse regions, habitats, and taxa. iNaturalist also provides means for users to annotate information about images beyond an identification, including leafing, flowering, and fruiting phenology [18]. However, while millions of images on iNaturalist have such annotations, the vast majority of iNaturalist images are not annotated with phenology information and annotation pace is far slower than new observation uploads.

A key next step for unlocking phenology data resources at scale is using computer vision to augment community science efforts, scaling production of phenology data availability by orders of magnitude. Machine learning-based computer vision methods provide the means to generalize associations between image features and phenological stages such as flowering and fruiting. By training machine learning models on human-annotated images and applying a tuned model to the tens of millions of iNaturalist field images without annotations, researchers can unlock vast phenological information. This human-machine collaboration amplifies the impact of community science efforts, harnessing the strengths of both human expertise and machine efficiency to dramatically enhance scale, accuracy and completeness of phenological datasets.

Community science and machine learning together provide a powerful means to scale the assembly of phenology information, but do not alone solve quality control issues. Using computer vision methods on community-collected images is challenging given both varying quality and complexity of field images, and inherent difficulties with detection that vary across plant taxa. Beyond developing a performant modeling framework for annotation, careful vetting and filtering of results is needed to assure data quality reaches standards usable for downstream research. These steps and resulting information about data quality must also be fully articulated to users to build understanding and trust of the resulting data. As well, because community science resources are growing, both in terms of digital occurrence data records and in human annotations, pipelines need to be developed using cycles of continuous improvement and integration.

Here we describe PhenoVision, a new machine learning modeling pipeline utilizing iNaturalist occurrence records and associated field images of plants to enable scalability of phenology data via the automation of flower and fruit annotations. In particular, we cover the challenges and pitfalls working with community science images at scale, especially how careful decision making and tuning of post-model outputs helps to significantly decrease false positive error rates while retaining the vast majority of high quality phenology annotations. We discuss critical and strategic choices made to ensure stakeholders can understand, trust, and utilize our outputs for downstream phenology research. We also provide data and tools for access to PhenoVision data. We close by discussing next steps and plans to integrate PhenoVision outputs with data from in situ phenology monitoring programs to extend this initial effort at enhancing global phenology data resources.

## Methods

### Quality of user-contributed phenology annotations

The first step in the development of our machine learning model was curating a reliable and extensive training dataset of human-annotated images. The iNaturalist platform has a feature allowing users to annotate images with phenological information, with categories including flowers, flower buds, fruits or seeds, and no flowers or fruits. To ensure these annotations were accurate enough to use as training data, we manually validated a random sample of 2000 images in each of the categories except flower buds. Our validation results confirmed that iNaturalist user phenology annotations for flowers and fruits are highly accurate (see Results for more detail), and therefore usable as training data for our model. For the first phase of PhenoVision, we converted annotations into labels by coding them into a vector of two binary values: one indicating the presence (1) or absence (0) of fruit, and the other indicating the presence (1) or absence (0) of flowers. It is important to note iNaturalist only has absence reporting in cases where no flowers or fruits are present. Because of this, in cases where flowers were reported present but fruits aren’t, or vice versa, we assumed absence of the other trait. We discuss the impact of this issue with human annotations below. After curating annotations, we then downloaded all images with phenology annotation from iNaturalist as of March, 2024, using iNaturalist’s open data repository on AWS (https://registry.opendata.aws/inaturalist-open-data/). This gave us a training dataset of 1,535,930 research-grade images with associated annotated phenology data.

#### Training PhenoVision

We developed a machine learning model, PhenoVision, for detecting the presence of fruits and flowers in community-collected photos of plants. The model was trained on a large dataset of labeled images described in the above section. The neural network architecture of PhenoVision was based on a Vision Transformer (ViT) [19], specifically the ViT-Large variant, fine-tuned using a Masked Autoencoder (MAE) approach [20]. The ViT architecture, originally proposed by Dosovitskiy et al. [19], applies the transformer model, which has been highly successful in natural language processing, to computer vision tasks. In a ViT, an image is divided into fixed-size, linearly embedded patches, combined with position embeddings, and fed into a standard transformer encoder. This approach allows the model to attend to different parts of the image simultaneously, capturing both local and global features effectively. The MAE fine-tuning approach, introduced by He et al. [20], involves masking a significant portion of the input image patches (e.g., 75%) and training the model to reconstruct the missing pixels. This self-supervised pre-training technique has been shown to improve the ability of vision models to learn meaningful visual representations, enhancing performance on downstream tasks.

As a starting point, we utilized a pre-trained MAE model fine-tuned by Xu et al. [21] on the 2022 PlantCLEF competition dataset (https://www.imageclef.org/PlantCLEF2022), which comprises 2.9 million plant images labeled with species names, aggregated from various databases including iNaturalist and Pl@ntNet. Xu et al. [21] then additionally fine-tuned this pre-trained MAE on the task of classifying plants to species level. We reasoned that this pre-training would provide a useful inductive bias for our task, as plant species classification often relies heavily on the appearance of reproductive structures such as flowers and fruits, at least for human classifiers.

To adapt this pre-trained model for our specific task, we first discarded the final classification layer of the PlantCLEF-trained ViT. We then appended a new linear neural network layer with randomly initialized weights, which transformed the 1024-dimensional ‘latent’ representation of the masked autoencoder model into a 2-dimensional vector. This output was passed through a sigmoid activation function, ensuring final outputs consisted of two values ranging between 0 and 1, representing probability of fruit and flower presence, respectively.

To ensure robust model evaluation and prevent overfitting, we split our dataset into training, validation, and testing sets using a 60%, 20%, 20% split, respectively. This splitting was stratified by genus, flower presence, and fruit presence to maintain even proportions across combinations of these categories in each subset. The stratification strategy helped to ensure that each subset was representative of the overall dataset, reducing the risk of bias in model training and evaluation. This process resulted in training, validation, and testing sets of 921,720, 307,291, and 306,919 images respectively. We trained the model on the training set, while using the validation set to monitor model accuracy during training. This approach allowed us to detect and prevent overfitting, a common issue in machine learning where a model performs well on training data but fails to generalize to new, unseen data. The validation set was also used to determine an early stopping point for training, which occurred when the accuracy on the validation data no longer improved. Finally, we evaluated the model’s ultimate performance on the held-out test set, which provides an unbiased estimate of the model’s generalization ability. This three-way split strategy is standard machine learning practice and is important for several reasons. First, it allows for unbiased evaluation of the final model on data it has never seen during training or hyperparameter tuning. Second, it helps in detecting overfitting early, as performance on the validation set will typically plateau or decrease when the model begins to overfit. Third, it provides a more realistic assessment of how the model will perform on new, unseen data in real-world applications. By adhering to this rigorous evaluation approach, we can have greater confidence in the robustness and generalizability of the PhenoVision model.

The model was trained using stochastic gradient descent on batches of our training data. The model fit was optimized using binary cross-entropy as the loss function. During training, we optimized weights of the new neural network head while simultaneously fine-tuning the original weights of the ViT model. This approach, known as end-to-end fine-tuning, allowed the entire model to adapt to our specific task while retaining the beneficial pre-trained features from the PlantCLEF task. The model was trained using Python, with the pytorch [22], timm [23], and transformers [24] libraries, which were efficiently combined with R pre- and post-processing steps through the R package reticulate [25]. This integration allowed us to leverage the strengths of both programming environments, utilizing Python’s robust deep learning libraries and R’s data manipulation capabilities. We used a batch size of 2056 images for training. The model was trained on a single NVIDIA A100 GPU, with 8 CPUs dedicated to preprocessing and augmenting images and feeding them to the GPU in real-time. This hardware configuration allowed for efficient training of our large-scale model on the extensive dataset.

#### Post-training tuning of PhenoVision

We strategically opted to both reduce noise due to challenges with problematic images on iNaturalist, and to only focus on producing outputs for cases where flowers and fruits are detected. An overall objective of the effort is to determine accurately whether a flower or fruit is present at the whole plant level, allowing for robust downstream analysis based on known presences. Since images rarely show a whole plant, absence in an image does not guarantee absence on the whole plant at the time the image was taken. Since presence in an image can be taken as evidence for presence on the plant, but absence in an image cannot be taken as evidence for absence on the plant, we decided to prioritize accuracy of presence detection over accuracy of absence detection, in order to maximize performance on reliable evidence. Given these decisions, we developed a post-model calibration tuning pipeline that attempted to reduce noise from challenging images and to prioritize high accuracy of positive detections whilst minimizing data loss. To do so, we tuned the values of two important parameters for processing the output of the model after it had been trained: the ***detection threshold value***, and the ***uncertainty buffer zone***. The tuning was based on the held-out validation dataset to ensure independence from the training data, and was conducted with the R package {probably} [26].

The detection threshold value determines when a model output is considered a detection. A common threshold of 0.5 is often suboptimal, failing to maximize model evaluation metrics or accuracy based on validation datasets. We independently tuned this threshold for flowers and fruits due to differing class imbalances. Our methodology focused on optimizing Positive Predictive Value (PPV), Negative Predictive Value (NPV), True Positive Rate (TPR), and True Negative Rate (TNR) across various thresholds (Figure 1), using our independent validation dataset. We prioritized metrics related to positive instances: PPV to ensure model detections represented true instances, setting a minimum requirement of 0.9, while balancing it against TPR to minimize data loss.

**Figure 1.**
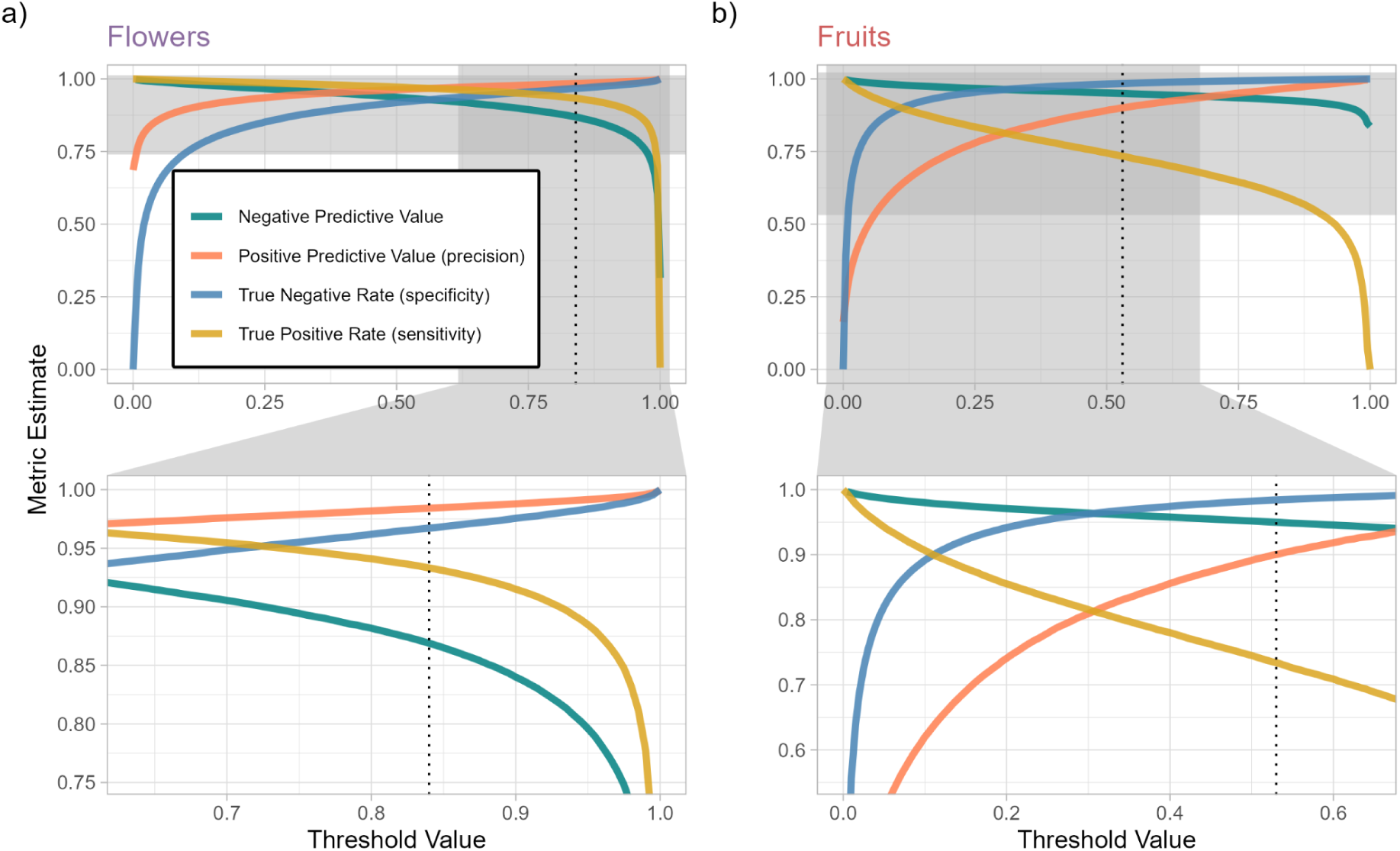
The panels show the relationship between model evaluation metrics and decision threshold values for flower (a) and fruit (b) detection in a binary classification model. The top row of panels show the behavior across the entire range of threshold values and metrics whereas the bottom panel shows a subset of the range of thresholds and metrics in order to highlight the behavior in the region of the chosen threshold. The dotted vertical line indicates the chosen threshold value, selected to balance Positive Predictive Value with other metrics, helping reduce false positive predictions, while maintaining acceptable levels of other metrics.

Once a detection threshold value had been determined, we optimized the value of a buffer zone around the threshold within which we would consider predictions as uncertain. It stands to reason that if a threshold is set to separate detections from non-detections, that values falling close to this threshold could represent ambiguous or uncertain predictions. We determined empirically how accuracy of the model on validation data changed around our chosen detection thresholds for fruits and flowers separately. In both cases we were able to determine a zone around the threshold where accuracy fell significantly. We decided to determine the borders of the uncertainty buffer zone by setting them where the accuracy first recovers to 75% on either side of the detection threshold.

#### Validating PhenoVision

Post model training, we performed human validation of model outputs. We manually selected and used our trained PhenoVision model to label 3,995 images for flowers and 1000 for fruits, and evaluated the model accuracy in each of these categories: detected-certain, detected-uncertain, undetected-certain, undetected-uncertain. During this step, we noticed that a relatively small, but consequential percentage of images were problematic for the purposes of phenology annotation in some way. These issues include problems intrinsic to the plants such as inconspicuous fruits or flowers, or fruits and flowers that cannot be distinguished from one another. There are also problems with image quality that are more extrinsic and due to the way a photograph was taken, such as images lacking a clear focal plant or images where the focal plant is too far away to see any features needed to assess phenology. To quantify the frequency of the most common extrinsic problematic field photographs, we manually scored a random sample of 500 iNaturalist images for presence/absence of a focal plant and presence/absence of a distance issue.

#### Further quality assessments

We tested overall model performance in three ways. First, we determined model performance using a 20% test holdout focusing on three metrics: Accuracy, True Skill Statistics, and F1 scores. Accuracy for detected and not detected is equivalent to the Positive Predictive Value (PPV) and Negative Predictive Value (NPV) respectively. The True Skill Statistic (TSS, also known as Youden’s J Index), is an overall model metric that incorporates both the True Positive Rate and the True Negative Rate, whereas the F1 Score, also an overall model metric, incorporates both precision and recall of the model. Besides prediction probability, a value between 0 to 1 representing a scaled probability of a presence or absence of a trait, we also defined and scored three other metrics useful for assessing quality of PhenoVision phenological labels. The first is “proportion_certainty_family”, the number of low certainty labels divided by the total number of machine-labeled iNaturalist observations in the taxonomic family. This is a metric of annotation success for observations in that family. We expect some families to be much more challenging to label than others given some groups have inconspicuous or difficult to differentiate flowers and fruits. The second is “accuracy_family”, a metric of per-family accuracy for the labels of all records in that family (certain or uncertain). This metric leverages both test and validation datasets, representing labels on >600,000 images. The last metric is “accuracy_excluding_certainty_family”, a metric of accuracy for the labels of only high certainty records.

#### Labeling Millions of unannotated iNatural images

We built a custom downloader script for automating download of 53,535,189 images from a special-built AWS S3 bucket (https://registry.opendata.aws/inaturalist-open-data/) which contains angiosperms images with creative-common licenses that permit non-commercial reuse.

These images were then stored on the University of Florida’s HiperGator main high-performance parallel filesystem, in order to assure performant labeling using the trained PhenoVision classifier. Post-classification, these images will be moved to archival storage since we expect that models will be re-trained and run again given both increasing label data being collected and to capture more phenology phases.

We created a custom pipeline using the R package {targets}, which implements a powerful make-like workflow with automatic dependency detection [27]. The pipeline does the following: 1) splits the images into batches that can be processes on a single A100 GPU, 2) applies a standard image center cropping preprocessing step, 3) generates Phenovision model output for each image, 4) applies our tuned detection threshold to generate detections or non-detections, 5) uses out tuned uncertainty buffer zone to label each detection as certain or uncertain, and finally 6) filters out non-detections and uncertain detections. Using this pipeline we ensure that the machine labeling procedure can be interrupted and restarted, or rerun using new models or datasets whilst making sure the same combination of model and image will never need to be run more than once, reducing unnecessary computation and energy use. After using this pipeline on the current set of downloaded unannotated iNaturalist images, we had a final dataset of 30,199,391 phenology records, 25,276,056 for presence of flowers, and 4,923,335 for fruits.

#### Ingesting PhenoVision data in a standardized format

After completing labeling with PhenoVision, we developed and implemented a standardized format for reporting outputs that includes key information about each annotated record needed for phenology estimation, including taxon name, latitude and longitude, day of year, year collected, iNaturalist URL link, and the iNaturalist record associated with that image. The output format also includes the trait being predicted (e.g. “flower” or “fruit”), the prediction class (“detected” or “not detected”), and a model uri, providing a means to link all trait labels to a particular model run. We then developed an ingest pipeline and application to provide access to these data. We first filtered out all reports of flowers or fruits that were “uncertain”, and also removed the observations where the outcome was “non-detection”. We did so because “uncertain” labels were validated to have high error rates (see below), and removing these only led to a marginal (∼7%) decrease in overall number of records. We removed non-detections, for three reasons: 1) our approach produced higher quality for detections, 2) non-detections are not evidence of whole plant absence of flower or fruit; 3) iNaturalist data is best suited for presence-only approaches as they are incidentally collected.

Finally, we mapped the output file with key terms in the Plant Phenology Ontology (PPO) [28] and utilized an existing pipeline to produce an ontology mapped final dataset. The mapping to the PPO provides a means for integrating iNaturalist data with other data sources such as in situ observatory data from the USA National Phenology Network [28] and herbarium sheets annotated for phenology [29]. The pipeline creates a reasoning step that creates new information, e.g. that “plant reproductive structures” must be present if either flowers or fruits are present. The end result was a finalized dataset of plant phenology labels, which we ultimately make available for community use (see Results).

#### Evaluating post-PhenoVision coverage

We determined how much machine labeling of flower and fruit presence adds to the existing corpus of annotations produced by public participants from iNaturalist. We focused on two different views of coverage gains. The first is a visualization of gain in coverage spatially. We used R package {sf} [30] to create a 100 km by 100 km equal-area cells map globally and determined both the absolute number of new records added by PhenoVision, and the percent gain of records. Percent gain is simply the number of records generated by PhenoVision divided by the total number of annotated records (computer or human) available in a grid cell. The second visualization provides a view of the gain in phenology records phylogenetically. For each genus, we determined the percent of total from human and computer labels and plotted those percentages per genus on an angiosperm phylogeny. We used R packages {rtrees} [31], {ggtree} [32], and {ggtreeExtra} [33] for assembling a synthesis phylogeny and plotting ratios across each generic-level tip in the tree.

## Results

### Quality of user-contributed phenology annotations on iNaturalist

We found user-contributed phenology annotations on iNaturalist to be highly accurate for flower presence, fruit presence, and flower and fruit absence (Table 1). Images annotated with flower presence were 97.5% accurate and fruit presence were 92.3% accurate (Table 1). Images annotated with both flower and fruit absence were 97.5% accurate for flowers and 96.7% accurate for fruits. We marked a small subset of images (1%-5% depending on category) as unknown, meaning we were unable to determine fruit or flower presence from the image (Table 1). This unknown category included photographs where focal plants are hard to distinguish as well as plants in families where reproductive structures are challenging to distinguish from an image alone (e.g. Poaceae, the grasses). The high accuracy values for all three relevant phenology annotation categories allowed us to move forward with using iNaturalist user-contributed phenology annotations as training labels. Our final training dataset contained 1,535,930 images with associated phenology data from iNaturalist.

**Table 1.** Expert validation results for user-contributed phenology annotations on iNaturalist. The rows represent the status of a phenological stage according to an expert annotator, whereas the columns are the presence or absence of stages according to iNaturalist annotators, split by flower or fruit. Cells in bold represent the cases where the expert and iNaturalist annotators agreed on the phenological state, whereas unbolded cells are where they disagreed, or the expert could not determine the correct state from the image. iNaturalist has one category for absence (e.g. flowers and fruits absent) and expert annotation was for the target listed in parentheses for absences.

| Expert annotation | iNaturalist<br>flowers present | iNaturalist<br>fruits present | iNaturalist absent<br>(flowers) | iNaturalist<br>absent (fruits) |
| --- | --- | --- | --- | --- |
| Stage present | **97.5%** | **92.3%** | 0.7% | 1% |
| Stage absent | 1.2% | 2.6% | **97.5%** | **96.7%** |
| Unknown | 1.3% | 5.1% | 1.8% | 2.3% |

### Model Calibration and Tuning

We downloaded images and metadata, including phenology annotations, from an AWS image store (URL) made available by iNaturalist, that included only images with a CC-0, CC-BY, or CC-BY-NC license. Model calibration was performed as discussed in the Methods and here we focus on post-model tuning results. The model converged to an asymptotic minimum loss very quickly, achieving a high level of accuracy on validation data after only four epochs (four complete passes through the full dataset). This rapid convergence suggests that pretrained representations from the PlantCLEF task were indeed relevant to our phenological detection task.

As discussed in Methods, we determined two key post-calibration model tuning parameters, a ***detection threshold value***, and the ***uncertainty buffer zone***. Regarding the detection threshold value, Figure 1 shows results for how well key metrics performed across a range of threshold scores for classifying presences for flowers and fruits. For flowering, we set the threshold at 0.84, which maximized the True Skill Statistic (which itself is proportional to the mean of TPR and TNR). This yielded a high PPV while maintaining a strong TPR (Figure 1, Panel A). For fruiting, due to steeper trade-offs between PPV and TPR, we set the threshold at 0.53, where PPV equaled 0.9 and TPR was 0.75. This tuning process allowed us to optimize model performance, addressing class imbalance issues and prioritizing accurate positive detections for both flowers and fruits.

Figure 2 demonstrates the relationship between model confidence, accuracy, and the number of images classified at different output levels, aiding in the selection of optimal thresholds for flower and fruit detection to maximize accuracy of positive detection whilst losing as little data as possible (uncertain images were discarded from the database). This result provides a means for determining a zone of uncertainty around the detection threshold value, shown in Figure 2 for both flowers and fruits. By excluding model outputs in the uncertainty zone in our final annotation dataset, we were able to substantially increase accuracy of the model whilst losing only a relatively small amount of data (approximately 7%, based on the validation dataset, for both fruits and flowers; Table 2).

**Figure 2.**
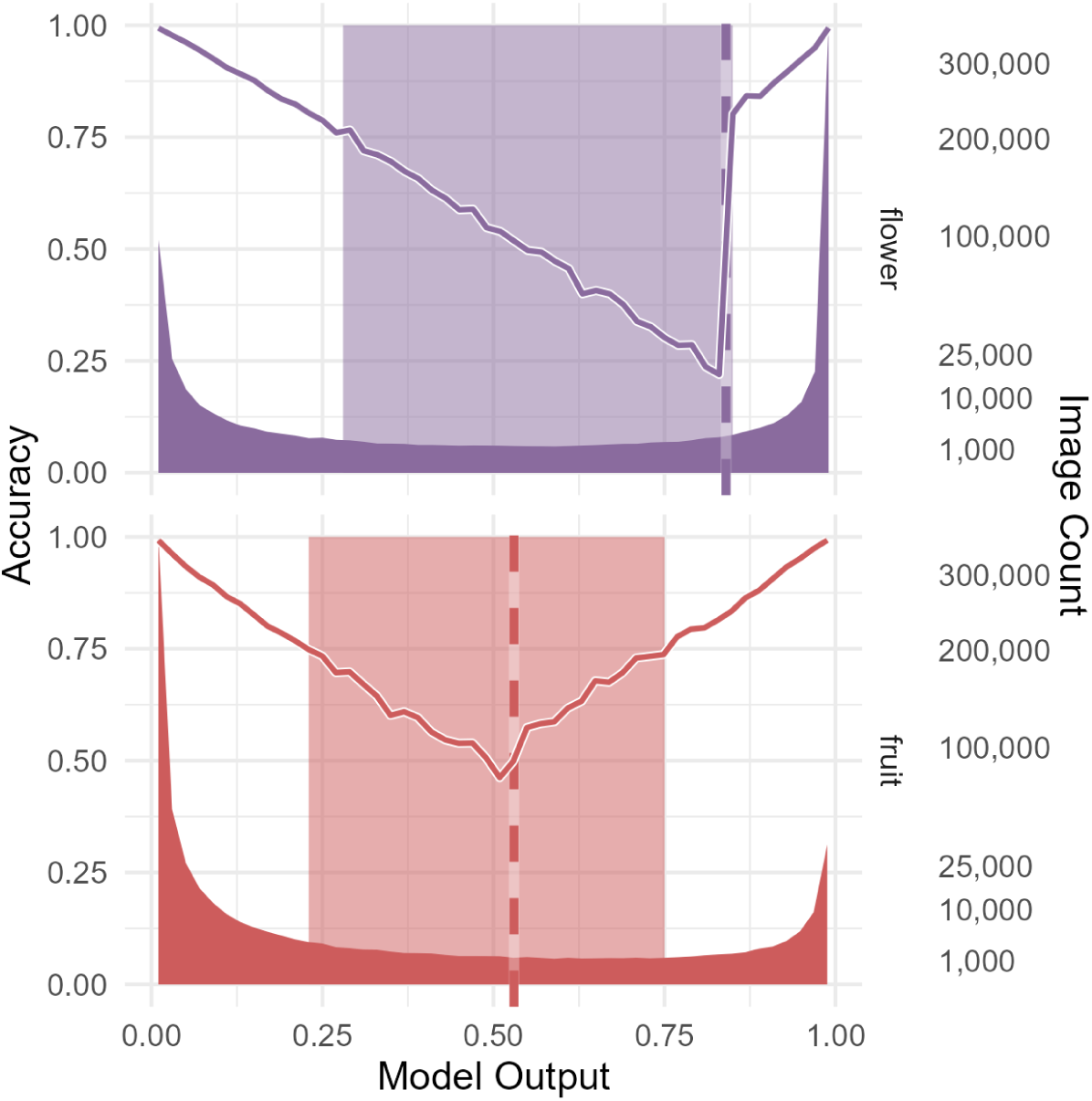
Distribution of model outputs and accuracy for flower (top) and fruit (bottom) detection. The x-axis represents the model output (probability of detection), while the left y-axis shows the accuracy, and the right y-axis (square root scale) shows the image count. The darker shaded areas represent the distribution of image counts and the lighter shaded areas highlight the range of outputs considered ‘uncertain’ (e.g. the buffer zone). The solid lines indicate the accuracy and the vertical dashed line represents the chosen decision threshold for each trait – any images with output values above this would be considered a detection, below a non-detection.

**Table 2.**
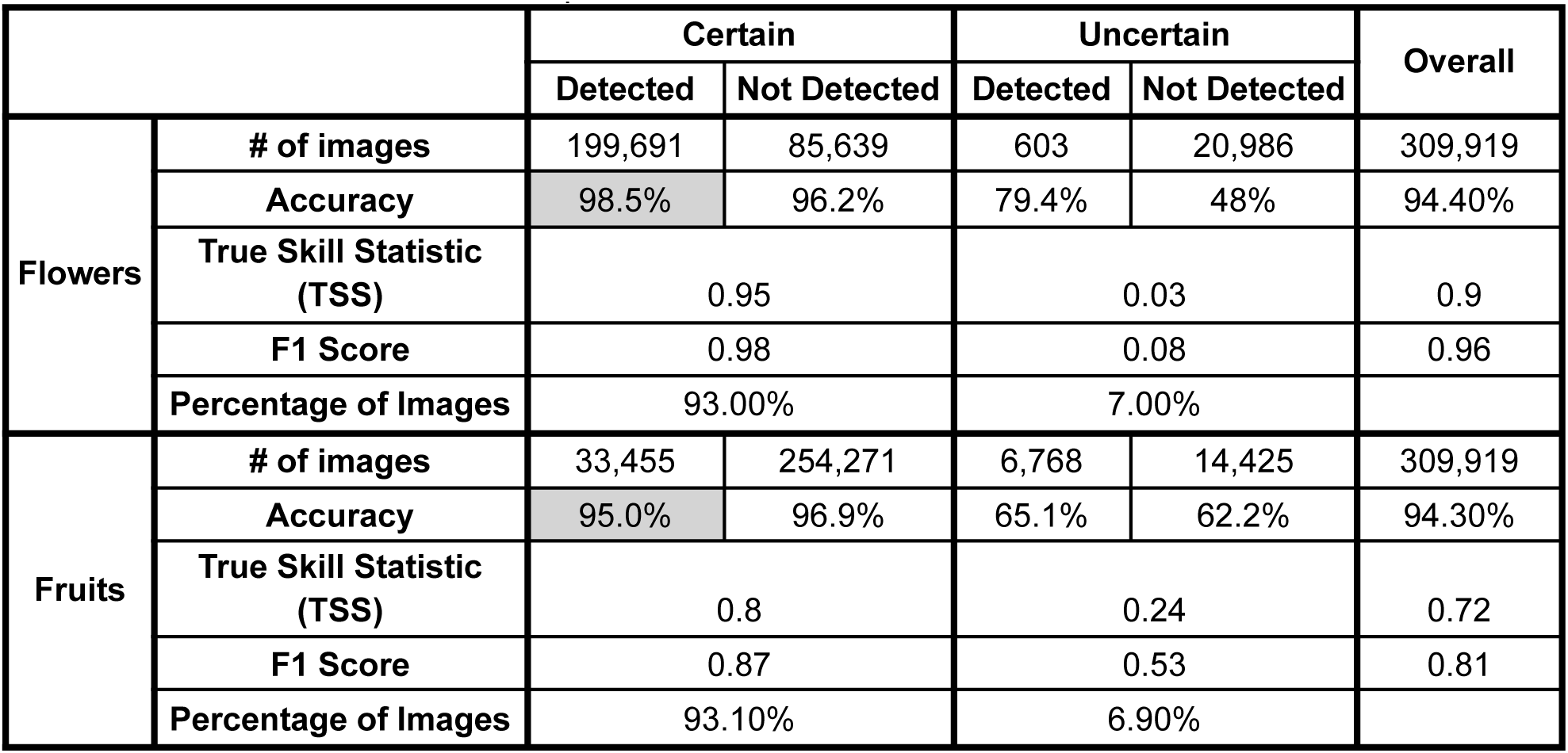
Validation result for model-produced flower and fruit annotation on held-out test dataset (20% of original human-annotated data). We show accuracy split by certainty and by whether the flower or fruit was detected by the model. Shaded cells represent accuracy scores for machine-labeled data that will be retained and included in the final data product.

|  |  | Certain |  | Uncertain |  | Overall |
| --- | --- | --- | --- | --- | --- | --- |
|  |  | Detected | Not Detected | Detected | Not Detected |  |
| Flowers | # of images | 199,691 | 85,639 | 603 | 20,986 | 309,919 |
|  | Accuracy | 98.5% | 96.2% | 79.4% | 48% | 94.40% |
|  | True Skill Statistic (TSS) | 0.95 |  | 0.03 |  | 0.9 |
|  | F1 Score | 0.98 |  | 0.08 |  | 0.96 |
|  | Percentage of Images | 93.00% |  | 7.00% |  |  |
| Fruits | # of images | 33,455 | 254,271 | 6,768 | 14,425 | 309,919 |
|  | Accuracy | 95.0% | 96.9% | 65.1% | 62.2% | 94.30% |
|  | True Skill Statistic (TSS) | 0.8 |  | 0.24 |  | 0.72 |
|  | F1 Score | 0.87 |  | 0.53 |  | 0.81 |
|  | Percentage of Images | 93.10% |  | 6.90% |  |  |

### Validating Model Performance

We tested the success of our models for classifying flowers and fruits present on iNaturalist images after post-model tuning described above. We used multiple methods for validation, including a 20% hold out of random test data and further human validation of new validation images. Table 3 summarizes the success of the models based on holding out 20% of the images with human annotations for testing. These results confirm that classifications that were in zones of uncertainty are significantly worse than those that were labeled as certain, especially for fruits. For both flowers and fruits, excluding uncertain model prediction led to a loss of about 7% of images from the dataset. In general, accuracy for the retained images are very high: 98.5% for flowers and 95.0% for fruits (Table 2).

**Table 3.** Validation results for PhenoVision-produced flower and fruit labels on 3995 random iNaturalist images for flowers and 1000 for fruits. The columns show the result from expert validation split into three categories. Scores in bold represent cases of agreement between the expert annotator and PhenoVision’s labels. Shaded cells represent agreement scores for labels that will be retained and included in the final data product.

|  | Flower |  |  |  | Fruit |  |  |  |
| --- | --- | --- | --- | --- | --- | --- | --- | --- |
|  | Detected |  | Not Detected |  | Detected |  | Not Detected |  |
| Expert annotation | Certain | Uncertain | Certain | Uncertain | Certain | Uncertain | Certain | Uncertain |
| Present | **98.6%** | **75.3%** | 2.1% | 56.4% | **90.4%** | **69.3%** | 15.4% | 49.1% |
| Absent | 0.4% | 12.5% | **82.9%** | **24.4%** | 3.4% | 18% | **77.6%** | **34.1%** |
| Unknown | 1% | 12.2% | 14% | 19.2% | 6.2% | 12.7% | 7% | 16.8% |

**Table 4.** The frequency of problematic images in iNaturalist and the resulting model decisions for these problematic images. Each cell represents a percentage of a representative sample of all iNaturalist images. Shaded cells represent the percentage of iNaturalist images that are problematic **and** will be included in the final data product.

| Flower |  |  |  | Fruit |  |  |  |
| --- | --- | --- | --- | --- | --- | --- | --- |
| 10.0% of images are problematic |  |  |  | 12.2% of images are problematic |  |  |  |
| Certain |  | Uncertain |  | Certain |  | Uncertain |  |
| 7.4% |  | 2.6% |  | 9.6% |  | 2.6% |  |
| Presence | Absence | Presence | Absence | Presence | Absence | Presence | Absence |
| **2.0%** | 5.4% | 0% | 2.6% | **1.6%** | 8.0% | 0.4% | 2.2% |

We also evaluated models by randomly selecting and manually annotating 3995 images for flowers and 1000 for fruits. Evaluation here was validation of correct phenology state via expert assessment. Since our final data product will contain presence-only data in the “certain” category, we focus on the present and certain accuracy value for both flowers and fruits. For flower presence, the “certain” model labels are 98.6% accurate, and for fruit presence this value is 90.4% (Table 3). The decision to focus on presence-only data is supported by the relatively poor performance for model non-detections on plants where human validation shows definitive absence (82.9% accuracy for flowers, 77.6% for fruit). Many images were scored by human annotators as “unknown”, cases where annotation is not possible for a variety of reasons, including poor image quality or taxa with inconspicuous flowers or fruits. Unknowns from human classifiers are far more common in the uncertain model category, especially for detections, with only 1% of certain flower presences are labeled as unknown. The rates are higher for certain detections of human-scored unknowns for fruits.

We further examined a separate random subset of 500 iNaturalist images to examine cases where iNaturalist images were scored as unknowns due to extrinsic photo issues, with a focus on better understanding the kinds of problematic images and their impact on classification success. We found that, out of all investigated iNaturalist images, 10% of images are extrinsically problematic for human flower detection and 12.2% of images are extrinsically problematic for human fruit detection (Table 3). However, in the final data product, which will contain presence-only “certain” data, only 2% of images are problematic for flowers and 1.6% for fruits. These percentages represent images that are classified by the model for phenology, but can’t be verified by human experts. Finally, model accuracy varies taxonomically, as some groups of plants have small or otherwise difficult to determine flowers or fruits. Supplemental Figures 1 and 2 show model accuracy by family, which demonstrates variability in accuracy, with some families such as Halogoracea demonstrating much lower accuracy than most families. We cover more about issues and challenges with machine annotation of certain groups in the Discussion.

### Model Errors

We looked at a random sample of images that were incorrectly scored by the model to gain an understanding of what situations might cause the model to make mistakes. There are two error types: false positive errors, which occur when the model reports a presence of a fruit or flower but our human validation data reports an absence, and false negative errors, which occur when the model reports an absence but our human validation data reports a presence. We focus primarily on sources of false positive errors, as these are the only errors which make it into our final, presence-only dataset. We found, anecdotally, for both flowers and fruits, that the majority of false positive errors are actually true positives, meaning the model annotation is correct and the human validation annotation is incorrect. For flowers, this phenomenon is almost entirely explained by an issue with incomplete iNaturalist phenology annotations (see “annotation quality” section of Methods). For fruits, we see this as well, but we also see more cases of incorrect human annotation. We note other causes of false positive errors for cases when the model was truly incorrect, including mistaking other reproductive structures for the reproductive structure of focus, scoring reproductive presence on the non-focal plant when the focal plant has an absence, and scoring a presence for an image where phenology is impossible to determine (e.g. image taken too far away) (Figure 3).

**Figure 3.**
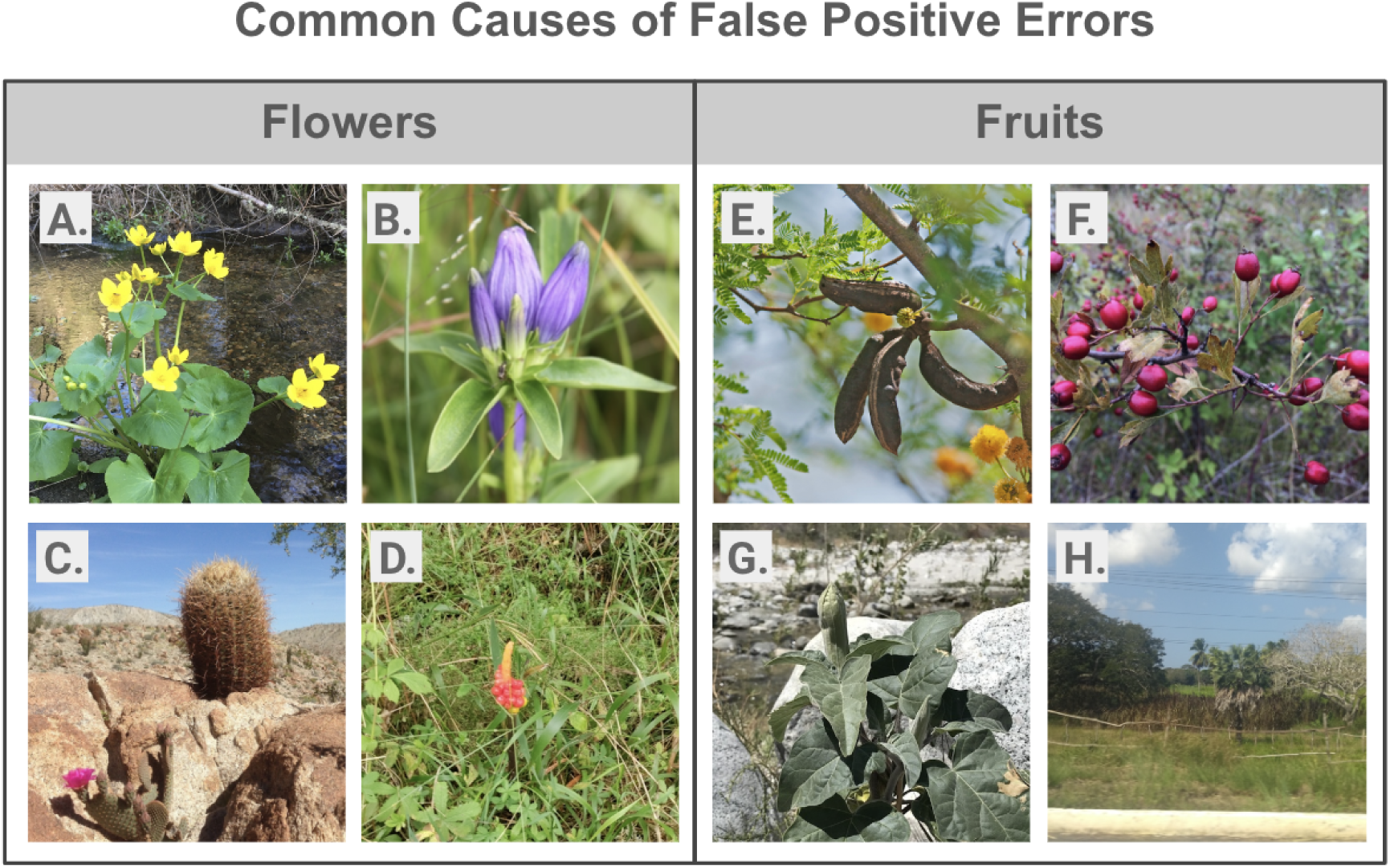
Examples of common reasons for false-positives, situations where the model reports a presence but our human validation data reports an absence, for both fruits and flowers. **A.** There is a flower in the image, e.g. the model is correct and the image is actually a true positive. This is the most common “error” for flowers and is due to incomplete annotations on iNaturalist. **B.** The model mistook unopened buds for flowers. **C.** The non-target plant has a flower, but the target plant, which is the species linked in the iNaturalist observation, does not. **D.** The model mistook fruits for flowers. **E.** There is a fruit in the image, along with a flower. The model is correct and this is a true positive. This is a common issue resulting from incomplete annotations on iNaturalist. **F.** There is a fruit in the image with no flower. The model is correct and this is a true positive. This situation is not an issue with an incomplete annotation, but demonstrates that the human annotator incorrectly scored for fruits. This issue is relatively common for fruits, but rare for flowers. **G.** The model mistook flower buds for fruits. **H.** The model annotated fruit presence for an image that was taken at a distance too far to tell if fruits are present or absent.

### Vastly Increased Coverage of Standardized Phenology Data Globally

We developed a standardized final dataset of 30,199,391 phenology records, 25,276,056 for presence of flowers, and 4,923,335 for fruits. These standardized records are Plant Phenology Ontology compliant and contain key data from iNaturalist records that support direct use for phenology analyses. In particular, each record contains information on day of year and location of the recording event, taxon recorded, along with key data about the annotation process such as reproductive structure (e.g. flower or fruit), classification outcome and probability score, quality metrics and model identifier. We ingested the final mapped dataset into a simple web application that provides filtered searching by date, taxon and phenology trait and classification outcome (https://phenobase.netlify.app/).

We next used these new data to determine how coverage increased both phylogenetically and spatially after adding to existing human annotation based data in iNaturalist Phylogenetically, PhenoVision generated phenology data for 119,340 species from 10,406 genera and 408 families. For the 7,409 genera with at least 10 flower records, the median and mean number of machine annotated flower records per genus was 184 and 3,276, respectively, with the maximum number of 357,608. For these genera, the minimal proportion of records that were generated by PhenoVision was 15% (genus Prosopis, which had 353 human annotated records and 63 machine annotated records) while the 25th percentile proportion jumped to 95% (Figure 4, panel A). The pattern is similar for fruits. For the 5,240 genera with at least 10 fruit records, the median and mean number of machine annotated fruit records was 85 and 910, respectively, and the maximum number was 111,791. For these genera, the minimal proportion of records that were generated by PhenoVision was 10.5% (genus Astrophytum, which had 51 human annotated records and 6 machine annotated records) while the 25^th^ percentile jumped to 93% (Figure 4, panel B). The small proportion of machine annotated records for some of these genera was a result of a small number of available research-grade photos from iNaturalist.

**Figure 4.**
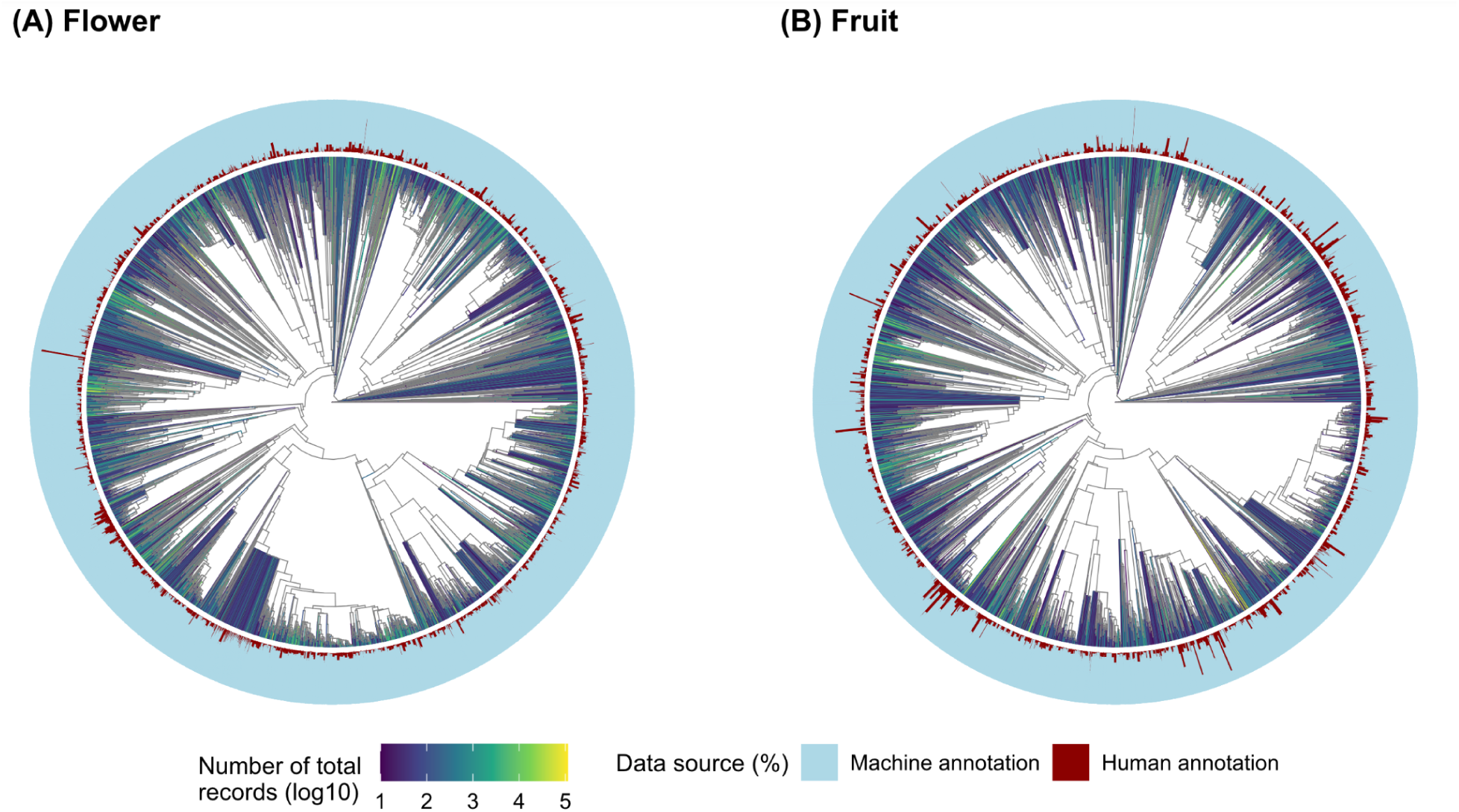
Machine labeled phenological data dramatically improves coverage of available phenology across the tree of angiosperms. Each tip in the phylogeny represents a genus with at least 10 total flower (A) or fruit (B) observations. Colors in the phylogeny branches represent the total number of phenology observations. Bars represented the proportion of human annotated flowering data (dark red) and the proportion of machine annotated flowering data (light blue) of 7409 genera for flower and of 5240 genera for fruit. For most genera, the vast majority of annotations are being generated by machine learning approaches.

Spatial coverage maps (Figure 5) show dramatic increases in data availability across the globe. PhenoVision generated phenology data for large areas where no phenology information previously existed in iNaturalist (3,798 and 4,147 new grid cells for flower and fruit, respectively; Fig. 5A), mostly from the global south and away from coastal and urban areas. In addition, PhenoVision largely increased the number of phenology records for areas with existing human annotated phenology data (Fig. 5 B-C), with many regions showing 1-4 orders of magnitude increase in number of records. Across all grid cells with existing human annotated phenology data, PhenoVision generated data now accounts for, on average, ∼90% of the total phenology data (Fig. 5C). For the 7,400 100km by 100km cells with less than 10 human annotations total before PhenoVision, there are now 1,577 cells that have at least 100 records annotated for flowering, and 236 cells with at least 500 records. This increase in available data should vastly increase the potential for conducting more advanced and more highly-powered analyses of phenology at the global scale. Still, many areas where biodiversity is richest are simply undersampled by iNaturalist, and remain gaps that are pernicious in phenology monitoring in general [10].

**Figure 5.**
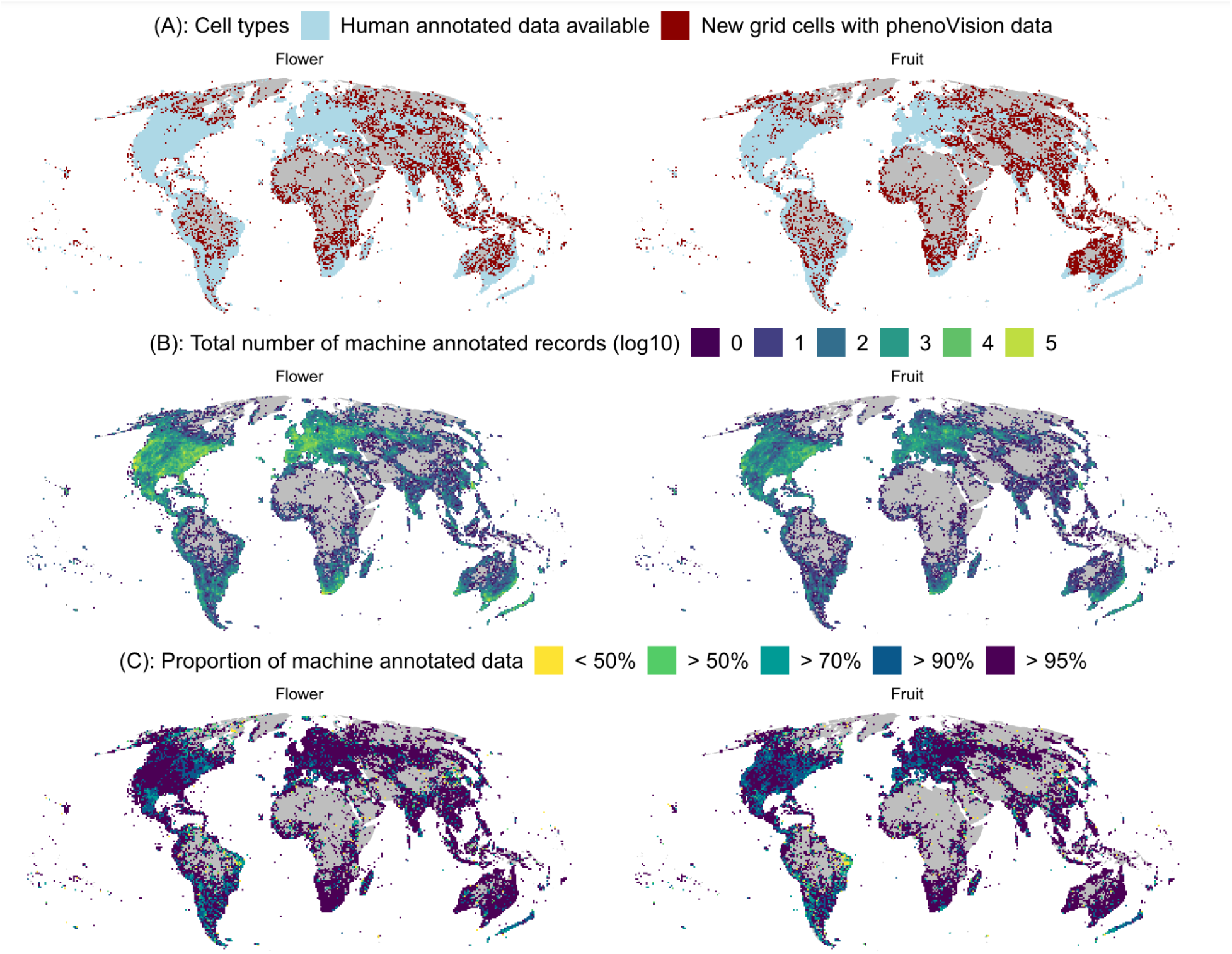
Global spatial-coverage of fruit and flower observation counts pre- and post-PhenoVision at 100km by 100km equal area grid cells level. (A) shows the spatial-distribution of grid cells with at least one manually-annotated flower and fruit presence record from iNaturalist (light blue) and new grid cells for which phenoVision provided data (dark red). (B) shows the spatial distribution of observation counts (log_10_-transformed) with flower and fruit labels generated by PhenoVision. (C) shows the spatial distribution of the proportion of phenology data generated by PhenoVision. Grey-colored grid cells contain no observation data.

## Discussion

Despite enormous progress in the application of machine learning in the biological sciences, full development of trustworthy community data and information resources built on ML approaches are still nascent. Here we focus on machine labeling of plant phenology and unveil PhenoVision, a new workflow which provides the means to annotate tens of millions of new phenology observations based on iNaturalist images and which is built to grow further. A key aspect of our work is a system design that is focused on four goals: 1) Highest quality outputs given the challenges with heterogeneous field images of plants; 2) Data that can be integrated with other sources of phenology information; 3) Ease of updating as new data becomes available; 4) Creating trustworthy, standardized data with details usable for assessing record-level quality. Below we discuss in more detail the lessons learned developing PhenoVision, in order to both support the best use of these phenology resources and to help others developing similar systems-based approaches. We close with key next steps and new frontiers with a focus on further automation, improving data integration and broadening how to extract new phenology traits from field images.

One of our key design decisions while developing the PhenoVision machine learning toolkit was not relying on traditional supervised machine vision models such such as VGG16 or ResNet which are based on convolutional neural networks, and typically pretrained on large heterogeneous databases of images such as imagenet. While known to be successful for tasks such as phenology detection in the past [34], recent work has revealed two particularly promising new strategies for producing high performing machine vision models: 1) Utilizing a transformer-based architecture known as a vision transformer (ViT), which are based on the success of similar architectures in Large Language Models, and 2) using a self-supervised learning as a form of model pretraining, where self-supervised learning can be conducted using unlabeled data to learn a task that might share important learning similarities to the labeling task of interest.

The masked autoencoder (MAE) approach suggested by [20] combines these two innovations by using a patch-based vision transformer architecture and a self-supervised task that aligns well with the architecture and creates powerful inductive biases for understanding image structure, specifically, the model is trained to reconstruct patches of the image based on the contents of other patches. Supervised learning is then used to fine-tune the model to a particular task, in this case phenology detection, by taking advantage of latent representations of the image data that the model pretraining has already learned. This is particularly powerful for scenarios like iNaturalist citizen science data, where we have a reasonable amount of labeled data, but a far more vast set of data that is unlabeled, which can be leveraged for the self-supervised learning stage. In the case of PhenoVision, this approach of combining self-supervised pre-training and a supervised classifier led to highly performant results, but this was just a key first step in our efforts to assure the highest quality computer annotations.

An essential component in ensuring trustworthy and high quality phenology labels was our iterative process of post-annotation human validation and resulting model parameter tuning and data filtering decisions. A key upfront choice made by our team was to only focus on images where flowers or fruits were present for several reasons. First, most of the images from iNaturalist only captured part of a plant individual, limiting ability to directly document that flowers or fruits were absent from the whole plant, even if the image lacks those structures. Because of this, absence data are often not utilized for downstream phenology research. Second, the model had a more difficult time with absences. We thus opted not to prioritize flower and fruit non-detections. This decision allowed us to focus on tuning quality for cases where reproductive structures were present, meaning we could prioritize high accuracy for fruit and flower detections.

Another key decision which ultimately led to higher-quality final outputs was to classify, and filter out, annotations where the model was uncertain. To do this we used plots of validation accuracy versus model classification scores (which range from 0 to 1) to empirically determine regions where models provide uncertain or equivocal versus more certain or unequivocal classifications for both fruits and flowers. The tuning step here represents a critical compromise between reducing the data volume of phenology annotations versus tuning for more quality. In our case, for both flowers and fruits, we were able to find clear cut off values that strongly improved quality (above 95% for fruits and 98% for flowers; Table 2) while only dropping ∼7% of data for either phenological phase.

Critically, filtering out annotations classified as “uncertain” often removes problematic images where it is difficult to determine phenological states of the focal, identified taxon, given the quality of the image. In particular, we identified two key classes of issues with detecting phenological states from community-collected field images, with some images having more than one kind of issue. The first class is when images contain non-target species in flower or fruit, but while relatively common, the issue is mitigated because it is relatively rare when focal taxa are not in flower or fruit when background taxa are, and because the model itself likely “learns” to tune out non-focal plant phenology states which contradict labeling of the focal plant. Supplemental Figure 3 (panel B) shows one example where this problem does happen. A second class of issues is images of landscape views, with a focal taxon far enough away where flower or fruit detection is not possible (Supplemental Figure 3, Panels C&D). This issue is more common for larger plants. Here we expect either uncertain model annotations or non-detections, even though in many cases these plants may indeed have flowers and fruits. Again, our decision to exclude non-detections is in part to avoid the possibility of significant false negatives.

Despite our efforts to tune models and post-process results to reduce noise, we still expect variation in the ability of the model to detect flowers and fruits. To acknowledge this, we produced further information that informs users about the taxa-specific quality of our final data outputs. Given that different angiosperm groups have varying difficulty in regards to phenology annotation, we provided users with two key metrics: proportional certainty per family (both before and after filtering uncertain records) and proportional family accuracy. These metrics give users a couple different views of whether records belonging to that taxonomic family are known to be easier or more difficult for the model to correctly label. We anticipate that users may decide to filter out certain challenging families for downstream use, e.g. Halogoraceae, which is known to have inconspicuous flowers and has the lowest family accuracy for flowers at ∼79%. An alternative in many cases is to focus on a broader phenological phase such as “reproductive structures present”, where overall accuracy may be higher, e.g. for Poaceae, where flowers and fruits can look similar. Finally, and critically, all model generated data maintains persistent URLs allowing users to immediately access both the photos annotated directly on iNaturalist and the records to which those photos are linked in cases where further validation is needed or wanted.

Although post-model validation and family-level certainty and accuracy metrics provide users with a deeper understanding of PhenoVision data quality, these evaluation methods are only as good as the annotation labeling data that inform them. Problems with detecting flowering and fruit phenology in certain taxa that are challenging for humans to annotate successfully, e.g. in many but not all grasses (Poaceae), may not be fully represented by these quality metrics. There are unavoidable errors in both the training and validation data, and error rates are likely higher for plant groups with small or difficult to distinguish reproductive structures. These errors can propagate into models and can lead to inflated validation statistics. We believe such issues are rare after careful scrutiny of model results, but further human validation before research use is likely valuable, especially for groups where flower and fruit detection is known to be challenging for human annotators. This point is not specific to computer-generated labels; the issue at root is the underlying quality of the annotations used to train the model (see Figure 3, panel A for an example).

### Challenges, Opportunities and Next Steps

PhenoVision has been built with expansion in mind, and this is a key design decision given the rapid growth of iNaturalist. We ingested and annotated all permissible images from iNaturalist prior to the end of March 2024. In just 6 months, as of the end of August 2024, about 10 million more images with permissive licenses can be annotated. While summer is the most active period for iNaturalist users, the rate of growth is impressive and our pipeline and toolkit is built to keep up with this pace, providing as close as possible to automated monthly or bi-monthly updates going forward. Many issues about phenological change relate to anomaly detections or impacts of extreme events [35] so providing near-real time resources is of significant value to the community. One key challenge is that not only are field photographs growing but so are annotations of phenological phases. These new annotations provide new fodder for re-training models, and a key design decision for PhenoVision has been tracking our machine learning model pipeline. Each model gets its own unique identifier, which is reported in the PhenoVision outputs we provide to users, and is linked to where the model is stored and easily downloadable from Hugging Face, allowing anyone to link models to model outputs. This means that retraining models or shifting to better modeling approaches later will allow us to produce new outputs while still maintaining older outputs. More work is needed across the machine learning community to specify standardized, lightweight model metadata that can link to these identifiers.

Beyond growth in annotations, there are significant opportunities for extending far past simple flower and fruit annotations. One area is to develop more subphases such as flower buds, open flowers, or senesced flowers. Further, detecting and counting individual flowers and their characteristics such as color or sex is rapidly becoming possible utilizing multiple different machine learning methods. Still, these next steps are challenging simply because they require being able to determine focal plants from background, and, while also true for our pipelines, the simplicity of our annotation targets is likely an advantage. For many questions, the simpler approach of documenting presence of a flower or fruit is sufficient for the research question being asked. These questions, while often phenological, may subtend into other areas such as interactions between plants and pollinators.

Another key area is developing annotations for other plant structures, in particular leaves, and this process is currently underway. As part of a collaboration with iNaturalist, iNaturalist has added leaf phenology annotations to its platform, including four categories: “breaking leaf buds”, “green leaves”, “colored leaves” and “no leaves”. We have already begun evaluating how well public participants are able to document these phenological phases, and already recognize that leaf annotations will have new and significant challenges when building automated pipelines. Most images that contain leaves of a focal taxon are correctly annotated, but background leaves are often present, far more than for flowering annotations. As well, some phenological phases, such as “breaking leaf buds” and “colored leaves” are challenging for iNaturalist users to document, and further may only make sense for a subset of taxa, requiring new downstream tuning.

A key next step in our process is integrating iNaturalist data with other data from monitoring programs, herbaria and other data sources to form a global central database of plant phenology, which we call “Phenobase” (https://phenobase.org/). The key to this integration is operationalizing the Plant Phenology Ontology [PPO; 28], which can greatly enhance both interoperability and findability across different systems with different reporting processes. All data generated from iNaturalist and PhenoVision were aligned to the PPO to assure such integration. iNaturalist phenology annotations, often a laborious process done manually by research teams, have already powered significant phenology findings [14,36] and that use continues to grow and expand. While more structured monitoring data on phenology is available in some countries and regions, often those programs are in developed regions and are focused on key taxa that are simple to monitor, and so are not comprehensive taxonomically. In sum, there are vast regions of the globe and vast numbers of taxa where direct data on phenology has been limited or non-existent. As we have shown in Figures 4 and 5, machine-annotated community science observations can significantly close global phenology data gaps.

We close by noting that despite the vast increase in available phenology data from our PhenoVision toolkit, these data are still as spatially and taxonomically biased as its underlying resources. While we are excited to see the availability of phenology in regions that are yet to be sampled by monitoring programs or made openly available to the community, gaps in coverage in all dimensions still exist [37]. Further, incidentally reported phenology data, while powerful for understanding phenology, still have limitations in terms of use for science and management compared to more semi-structured or structured monitoring [38]. Despite these limitations, continued integration of herbarium and in-situ phenology data resources (Phenobase) promise to provide the needed data basis for understanding global phenology trends in ways not previously imagined.

## Data and Model Availability

Full code for training, validating, and post-processing model is publicly available on Github: https://github.com/Phenobase/phenovision. The fully trained PhenoVision v1.0 model is available on Hugging Face (https://huggingface.co/phenobase/phenovision), and any future versions of the model will also be housed here to be freely used or reused by anyone in the community (MIT license). A dataset with all machine-labeled phenology records generated from this work is available on Zenodo (https://zenodo.org/uploads/13893483) and we have produced a demonstration application showcasing searching by taxon or phenology state here: https://phenobase.netlify.app/.

## Acknowledgements

This work is possible thanks to members of the iNaturalist community who take photographs of plants and upload them to iNaturalist (2.7 million people), add plant identifications (230,000 people), and annotate those observations with information about flower and fruit phenology (109,000 people). Our work has been funded by the National Science Foundation, in particular grants DBI2223508 to Daijiang Li and DBI2223512 to Robert Guralnick.

## SUPPLEMENTAL FIGURES

**Figure S1.**
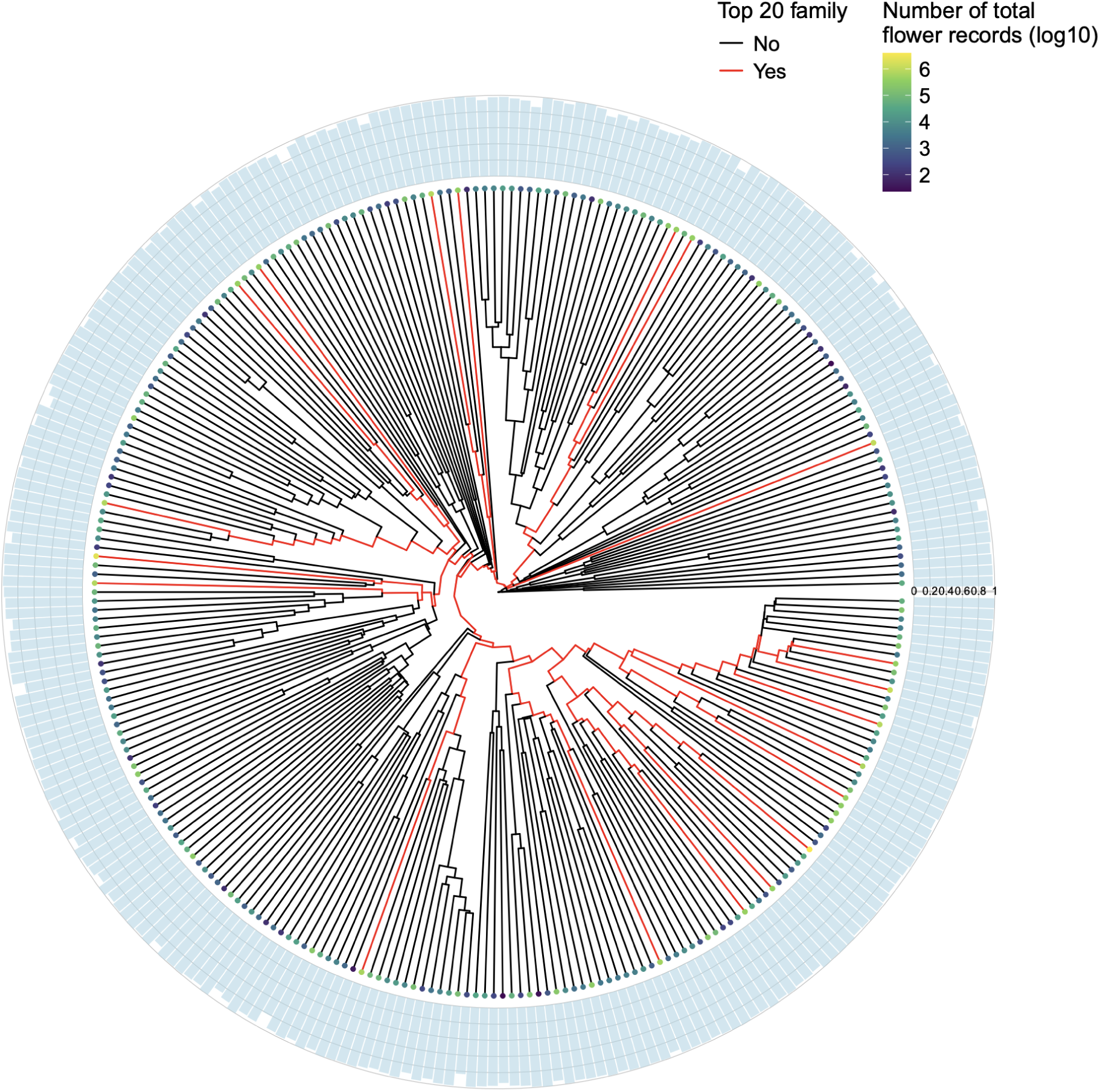
Accuracy of phenoVision for flower annotation at family level. We removed families with less than 10 flower records. The values plotted here ranged between 0.78 (Haloragaceae) and 1. The numbers of total flower records of families (ranged between 24 and over 3.8 million) were represented as colored dots at the tip top (at log-10 scale). The top 20 families with the most number of flower records (highlighted as red branches) have accuracy ranged between 0.969 to 0.995.

**Figure S2.**
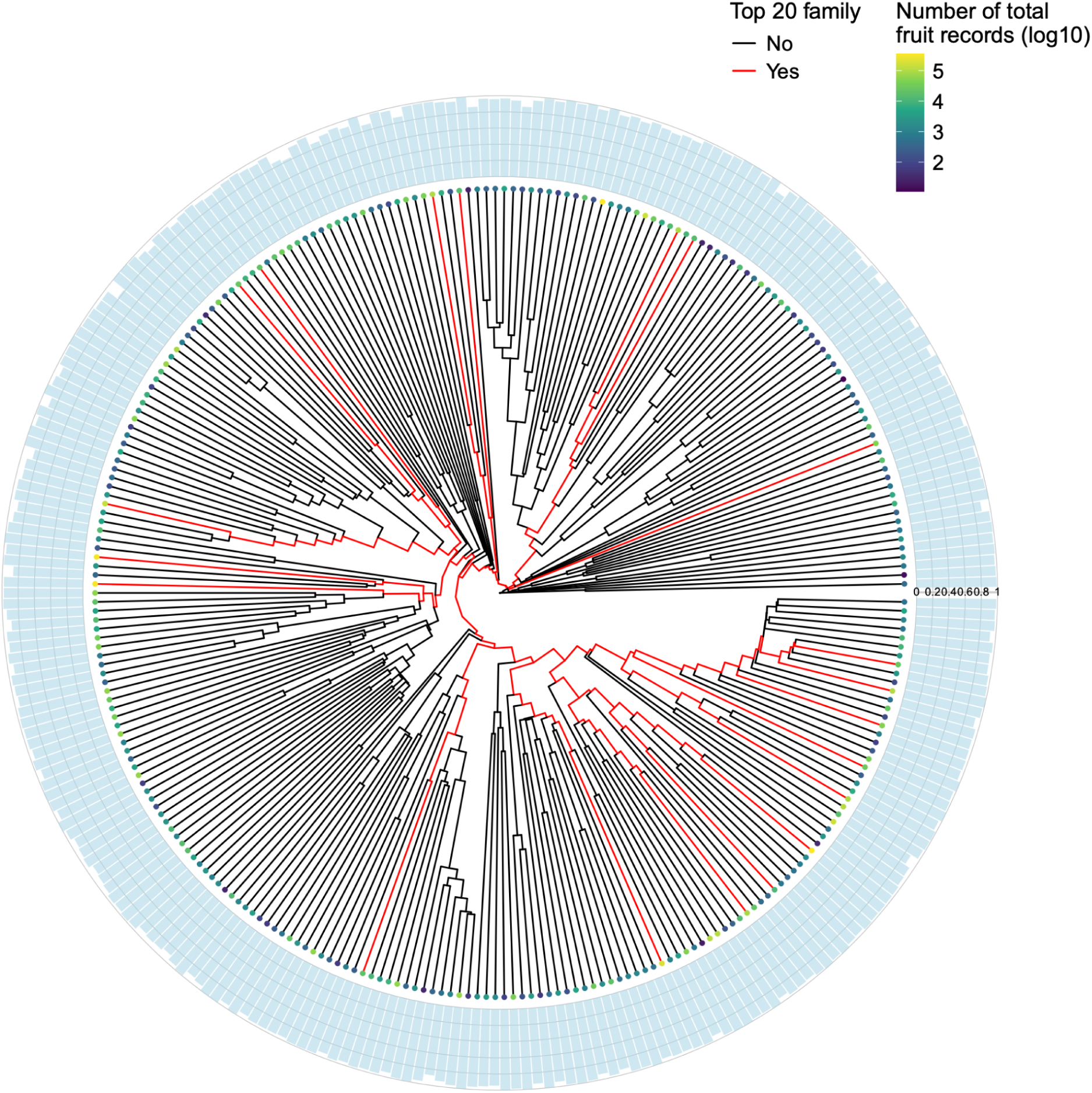
Accuracy of phenoVision for fruit annotation at family level. We removed families with less than 10 fruit records. The values plotted here ranged between 0.82 (Nitrariaceae) and 1. The numbers of total fruit records of families (ranged between 11 and over 355k) were represented as colored dots at the tip top (at log-10 scale). The top 20 families with the most number of fruit records (highlighted as red branches) have accracy ranged between 0.928 to 0.986.

**Figure S3.**
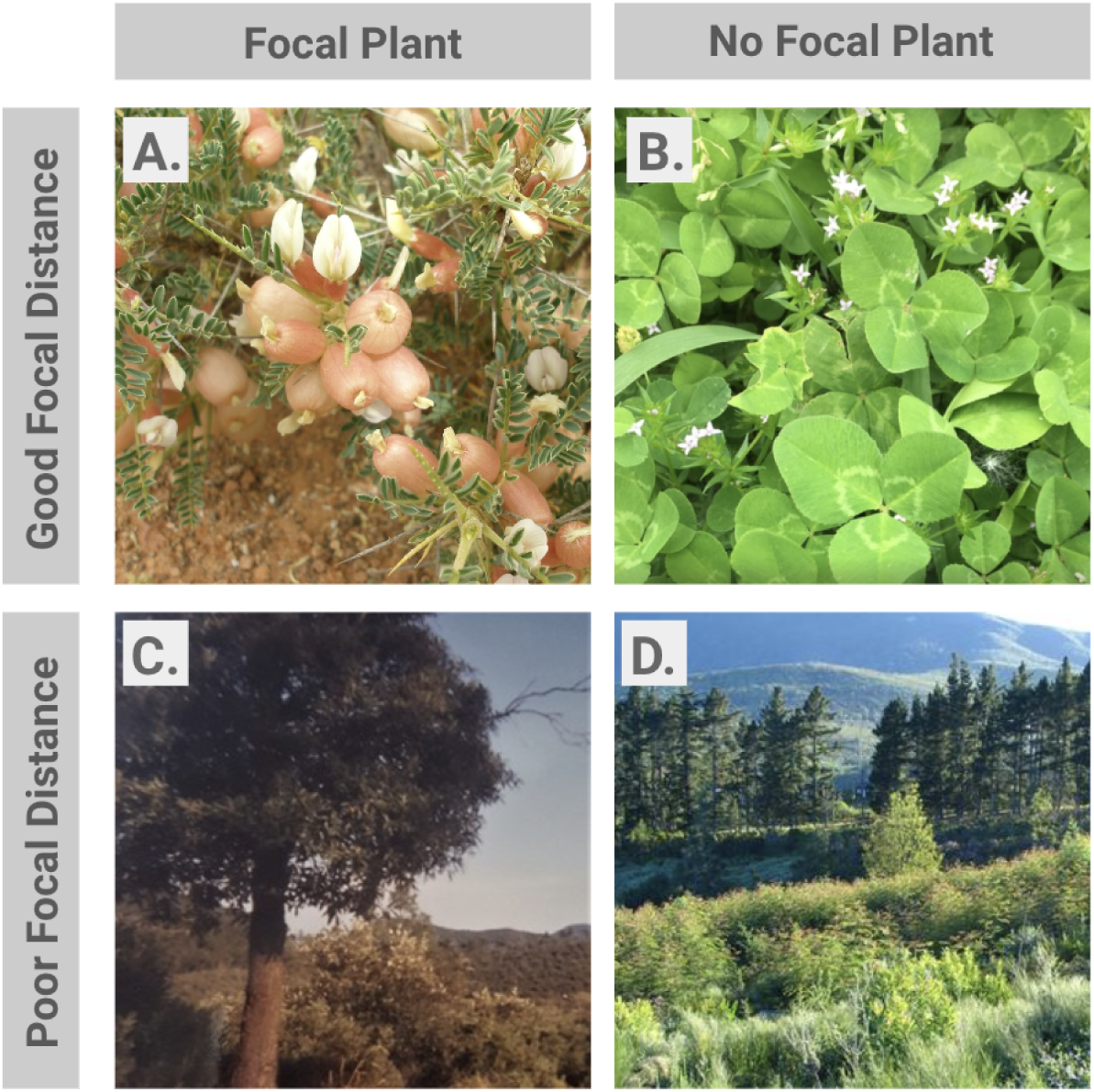
Examples of common iNaturalist image issues for flower and fruit recognition. A. Ideal iNaturalist image with no annotation issues. There is a clear focal plant and the photo is taken at a good focal distance, making flower and fruit recognition simple. B. iNaturalist image with no clear focal plant. Image taken at a good focal distance and reproductive structures can be recognized, however there are multiple plants that could be the focal species linked to the iNaturalist record. This becomes a problem when the multiple plants in the image have differing phenological states. C. iNaturalist image with a clear focal plant, but taken at a distance making reproductive structure recognition difficult. D. iNaturalist image with no obvious focal plant and taken at a distance making reproductive structure recognition difficult.

